# Transgenerational transmission of post-zygotic mutations suggests symmetric contribution of first two blastomeres to human germline

**DOI:** 10.1101/2024.06.18.599438

**Authors:** Yeongjun Jang, Livia Tomasini, Taejeong Bae, Anna Szekely, Flora M. Vaccarino, Alexej Abyzov

## Abstract

Little is known about the origin of germ cells in humans. We previously leveraged post-zygotic mutations to reconstruct zygote-rooted cell lineage ancestry trees in a phenotypically normal woman, termed NC0. Here, by sequencing the genome of her children and their father, we analyzed the transmission of early pre-gastrulation lineages and corresponding mutations across human generations. We found that the germline in NC0 is polyclonal and is founded by at least two cells likely descending from the two blastomeres arising from the first zygotic cleavage. Analyses of public data from several multi-children families and from 1,934 familial quads confirmed this finding in larger cohorts, revealing that known imbalances of up to 90:10 in early lineages allocation in somatic tissues are not reflected in transmission to offspring, establishing a fundamental difference in lineage allocation between the soma and the germline. Analyses of all the data consistently suggest that germline has a balanced 50:50 lineage allocation from the first two blastomeres.

## Introduction

Little is known about the origin of germ cells in humans. Human primordial germ cells (PGCs) are thought to be specified around 2 weeks post-fertilization^1,2^. While modeling germ-cells kinetics suggested that there are 2-3 founder germ cells (FGC)^3^, there is no direct evidence on the number of FGC and their ancestry, i.e., whether they are monoclonal or polyclonal. The major limitation to studying early lineages, including the germline, is ethical restrictions when working with human embryos. One way to overcome this limitation is to study cell lineages in parents using naturally occurring post-zygotic developmental mutations and then analyze their transmission to children. Post-zygotic mutations in the cells of a developing embryo start to accumulate from the very first cleavage and are acquired in almost every cell division^4-9^. As such, they represent records of development that are archived in the genome of every cell of a living person allowing to examine cell type and cell lineage allocations to all tissues including germline.

*De novo* variation in a child is defined as a mutation that happened in the germline of either parent. However, it was estimated that up to 6% of seemingly *de novo* variants in children can be detected at low frequency in parental blood, indicating that these “*de novo*” mutations are, in fact, early post-zygotic mutations of developmental origin in parents, arising prior to germline specification^4,10-13^. The standard trio approach commonly used to discover *de novo* variants by comparing blood-derived genomes of children to the blood-derived genomes of their parents is ineffective in finding early developmental mutations transmitted to the germline and then to children, since such mutations can be relatively frequent in parental blood and would be filtered out by this approach^4,10-13^. Using pedigrees with three generations (i.e., including the genome of grandparents in the analysis) allows discovering developmental mutations in parents^11^, yet often cannot distinguish true *de novo* from early post-zygotic mutations. This challenge can be addressed by the joint analysis of the genomes of large sibships (∼10 or more siblings), which are rare, where “*de novo*” mutations shared by multiple children represent early parental mosaic mutations transmitted to multiple cells in the germline. Such an analysis is inevitably indirect^10^. When applied to large Icelandic families this approach suggested the existence of 2 (or sometimes 3) FGCs^10^.

We previously leveraged post-zygotic mutations detected in clonal induced pluripotent stem cell (iPSC) lines to reconstruct zygote-rooted cell lineage ancestry trees in two living individuals)^5^. Briefly, we sequenced dozens of clonal lines derived from skin fibroblasts of each individual, discovered mutations present in the genome of founder cells for each line, analyzed shared mutations across lines, genotyped the mutation in tissues derived from the three germ layers, and reconstructed early developmental cell lineages starting from the likely first zygotic cleavage (**Fig. 1**). Lineages were assigned to the first zygotic division from the observations that descendants of the putative two blastomeres accounted for about 100% of cells in the tissues of the studied individuals, as measured by the frequency of corresponding mutations in blood, saliva, and urine. Similar results have also been obtained in post-mortem individuals^6-9^. A common finding emerging from these studies was that the two blastomeres resulting from the likely first cleavage of the zygote contribute asymmetrically to the soma. Such an asymmetry was as high as 90:10 in blood for both living individuals in our study, leading to our definition of dominant lineage (i.e., accounting for the majority of somatic cells) and recessive lineage (i.e., accounting for the minority of somatic cells)^5^. Here, by acquiring and sequencing the whole genome of the children of NC0, one of the two living individuals in our previous lineage study, and their biological father, we analyzed the transmission of early developmental mutations from a phenotypically normal woman to her children and elucidated the contribution of early developmental lineages to the germline. We next expanded our analyses to analyze transmission in a cohort of almost two thousand familial quads from the Simons Simple Collection.

**Figure 1.**
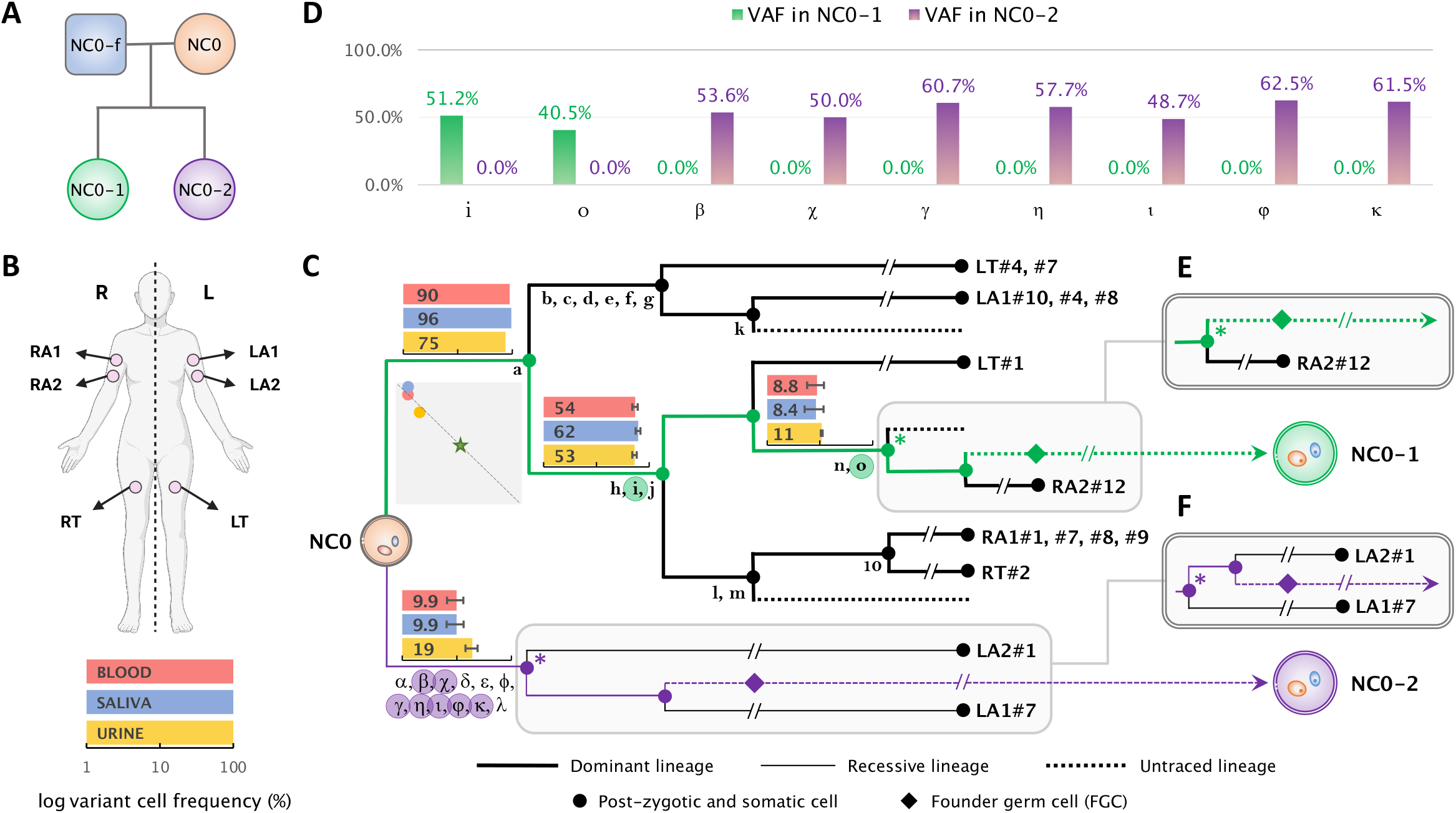
Early developmental cell lineages in the mother (NC0) and post-zygotic mutations tranmitted to the offspring. **(A)** Identifiers and relationship between 4 individuals (mother in orange; father in blue; two children in green and purple, respectvely) are shown in the pedigree for the sequenced family. **(B)** Body outline of NC0 shows location of biopsies used to derive iPSC lines from skin fibroblasts. **(C)** Cell lineage ancestry of NC0, which was originally reconstructed in our previous study^5^. Mutations shared by descendant cells are denoted by Latin and Greek letters including 4 indels (m, o, κ, λ). One branch has ten mutations and they are not denoted individually rather represented by a number. Mutations that were transmitted to NC0-1 and NC0-2 are highlighted in green and purple circles respectively. The corresponding cell lineages that bear the mutations transmitted to daughters are colored accordingly. For each branch in transmitted lineages, bar graphs show mean cell frequencies (corresponding percentages are indicated) of corresponding mutations in bulk blood (red), saliva (blue), and urine (yellow) in logarithmic scale. The squared plot shows correlations between the mutation cell frequencies (average across corresponding mutations; linear scale) in the three bulk samples for the dominant (y-axis: range 0 to 100%) and the recessive (x-axis: range 0 to 100%) lineages and their biased contribution to blood, saliva, urine (i.e., shifted away from the star which indicates equal contribution). Location of the dots along the dashed diagonal means that sum of frequencies for dominant and recessive lineage is 100%, i.e., the division is likely fully captured and all cells in analyzed tissues are progenies of the first two blastomeres. Diamonds depict founder germ cells (FGCs). The terminal of the branches represents the sampled clonal iPSC lines. **(D)** Variant Allele Frequencies (VAFs) in each child for all transmitted post-zygotic mosaic mutations in (C). **(E**,**F)** For the most recent ancestral cell (marked by “*”) traceable for the germline lineage that led to each offspring, alternative branches other than those shown in (C) are equally possible as shown in (E) for NC0-1 and (F) for NC0-2.

## Results

### Mutation transmission in the family of NC0

We found that 2 and 7 early post-zygotic developmental mutations traceable in the NC0 lineage tree originating from the dominant and recessive lineages, respectively, were transmitted to each child (NC0-1 and NC0-2) and were present in 100% of children’s blood cells, corresponding to a variant allele frequency (VAF) of 50% (**Fig. 1C and 1D; Supplementary Table 1; Methods**). In each lineage (dominant or recessive) mutations randomly originate on one of the two haplotypes. However, again randomly, only one haplotype will be inherited to the next generation, accounting for roughly 50% transmission of mutations in a lineage to the next generation. Consistent with that, each child had about half of the mutations transmitted from the inherited cell lineage (2 out of 5 for NC0-1 and 7 out of 12 for NC0-2) (**Fig. 1C**).

We next applied a common trio comparison approach to discover *de novo* mutations in each child by comparing their blood-derived genomes to those from the blood of the parents. Such a comparison discovered 84 and 94 (79 and 82 with more stringent filtering) candidate *de novo* variants in NC0-1 and NC0-2, respectively (**Methods**). Counts of *de novo* variants were consistent with the reported age of around 40 years old for both parents at the time of conception^14^. Importantly, only 1 to 4 (depending on the stringency of filtering) out of the 9 transmitted mutations could be identified as *de novo* using the trio strategy (**Methods**). This is because the trio comparison excludes from the “*de novo*” catalog all mutations detectable in parental blood, often missing variants (such as early developmental mutations) that are present in a significant proportion of cells in the blood of a parent (10%-90%; **Fig. 1C**). Consequently, the standard trio comparison^4,10,12,14,15^ mostly reveals *de novo* mutations that develop later within the parental germlines (and therefore absent in parental blood), and underestimates the number of mutations occurring prior to the specification of PGCs and transmitted to offspring; such that a child may have several (up to 6 in NC0-2) transmitted mutations not detected^14,15^. This observation also heights the unique power of direct lineage reconstruction and of the analysis of multi-children families (described below) for understanding mutation transmission across generations.

Each of the children of NC0 inherited early post-zygotic mutation arising from either the dominant or the recessive lineages in the mother. These observations proved that progenies from both dominant and recessive lineages in the soma are also FGCs in the germline of NC0. Therefore, the germline in NC0 consists of at least two clones (i.e., is polyclonal), each descending from one of the likely two blastomeres from the first zygotic cleavage.

### Mutation transmission in large families

We sought to confirm the observations in the family of NC0 using results from the Jonsson *et al*. study of transmitted mutations in families with multiple children^10^. In that study, personal haplotypes in children and parents were reconstructed using co-occurance of germline variants in the parental and children genomes. Transmitted mutations were inferred when a locus from the same haplotype was present in multiple children, carried a mutation in at last one child and did not carry the variant in at least one other child (**Fig. 2A** and **Supplementary Fig. 1**). Such transmitted mutations were often observed in parental blood and, thus, represent early post-zygotic mutations in parents. Because of relying on variant comparison across children and parents this approach is powered in revealing transmitted mutations when analyzing large families, with the power approaching saturation when there are 10 children in a family^10^.

**Figure 2.**
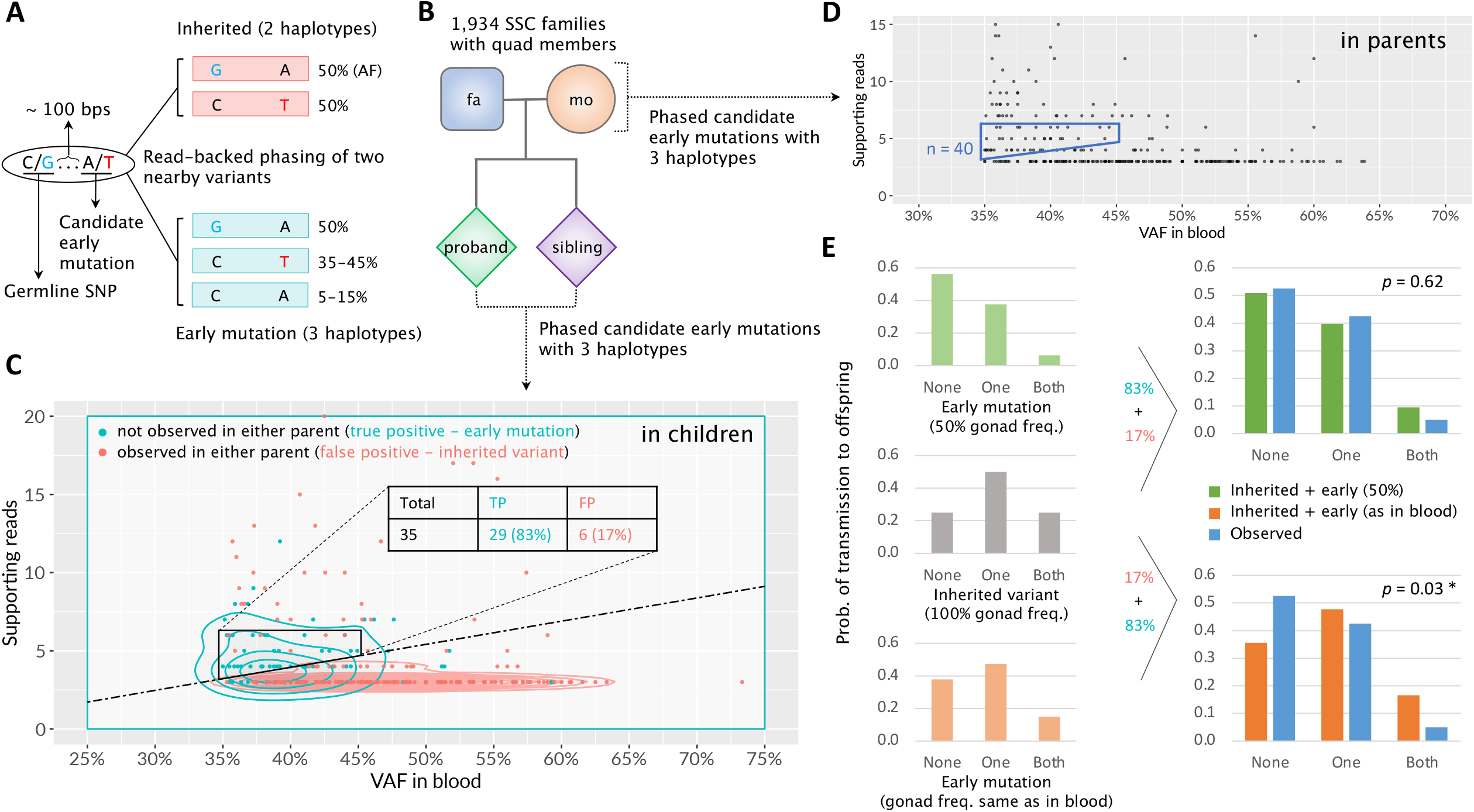
Early mosaic mutations and their transmission patterns to offspring in a cohort of 1,934 quartet families. Read-backed phasing approach to identify early mosaic mutations at high Variant Allele Frequencies (VAFs). A candidate early mutation with an alternative T allele (in red) is within <= 100 bps downstream of a heterozygous inherited SNP with alternative G allele (in cyan). If the candiate T allele is an inherited germline variant, we should observe only two haplotypes at a VAF of ∼50% for each. The presence of three haplotypes indicates that the candidate T allele is a post-zygotic mutation^11^. High VAF from 35% to 45% implies that the mutation occurred during the first zygotic division and marks the dominant cell lineage^5^. A pedigree structure of 1,934 familial quads in the Simons Simplex Collection^16^. **(C)** A scatter plot of quantitative characteristics for variants with evidance for 3 haplotypes in children. Each variant is represented by a dot with color corresponding to true post-zygotic murations (cyan) and false positives (red) that are observed in either parent and thus are inherited variants. Contour lines show density levels for the two categories of variants. Dashed line defines an approximate border demarkating the two distributions. Trapezoid by solid black lines defines the area to select early mutations in parents for the transmission analysis. **(D)** A scatter plot of characteristics for variants with evidance for 3 haplotypes in parents. Same as for children, the trapezoid delimited by solid blue lines defines the area to select early mutations for the transmission analysis. The analysis in children (panel C) allowed estimating that the false positives rate in this area is ∼17%. **(E)** Comparisons of expected and observed transmission rates of early post-zygotic mutations to offspring for two scenarios: (top) the mutation frequency is 50% in gonads (in green), i.e., dominant and recessive lineage contribute equally to gonads; (bottom) the mutations frequencies in gonads are the same as in the blood, i.e., VAFs are as in trapezoid in (D). There are three possible outcomes of mutation transmission: no transmission (denoted as “None”), transmission to only one child (denoted as “One”), and transmission to two children (denoted as “Both”). The expected distributions are calculated assuming 17% false positive rate from inherited variants. *χ*^2^ test with 2 degrees of freedom is used to assess the significance of the difference between expected (in green or orange) and observed (in blue) transmission distributions.

The analysis by Jonsson *et al*. of six large families with 17, 10, 9, 9, 9 and 9 children pointed to the existence of two (in a few cases of three) lineages in the germline of parents and their counterparts in the blood. For each parent in those families, we used transmitted mutation(s) with the highest variant allele frequency (VAF) in the blood to define the dominant lineage in the soma and estimate its frequency (**Methods**). In seven parents, we unambiguously identified the dominant lineage (contributing to >70% of cells) in the soma (**Table 1) (**see also **Figs. 2, S2**, and **S3 in Jonsson *et al***.^10^), similarly to what we observed in NC0. Strikingly, different from the soma, dominant and recessive lineages in those parents were roughly equally probable to be transmitted to children, and indeed, for all parents the frequency of transmitting the dominant lineage to children was always smaller than its frequency in the blood. Thus, the difference in the frequency of each lineage in the parental blood and its transmission to children was significant in four parents and, when combining analyses, across all parents and NC0 (combined *p*-value = 3.6 x 10^-17^). Jonsson *et al*. analyzed 251 families with multiple children and given the observed *p*-values (**Table 1**), random sampling of just one parent with such a difference is improbable. Thus, this analysis suggests that lineage allocation to germline and soma are drastically different, and the contributions of progenies from the first two blastomeres is asymmetric to soma but is symmetric to germline.

**Table 1.** Summary of frequencies and transmissions of dominant and recessive lineages in the current and in the Iceland pedigree study^10^. From the Iceland study we utilized data for seven parents from the largest families with clear existence (>70% VAF) of a dominant lineage in the blood. *P*-values by one-tailed binomial test reflect consistency in transmitting the dominant lineage to children relative to their frequency in the parental blood (*p1*) and consistency with respect to equal frequencies in the germline (*p2*). Last line sums children that inherit dominant and recessive lineage and provides combined *p*-value using Fisher’s method.

| Family | Parent | Cell frequency of the dominant lineage in the blood | # of children with lineage | | $p1$-value (as in blood) | $p2$-value (equal lineage contribution) |
| --- | --- | --- | --- | --- | --- | --- |
|  |  |  | dominant | other (recessive) |  |  |
| 17-children | Father | 93% | 8 | 9 | $6 \times 10^{-7}$ | 0.5 |
| | Mother | 94% | 6 | 11 | $3 \times 10^{-10}$ | 0.17 |
| 10-children | Mother | 72% | 5 | 5 | 0.12 | 0.62 |
| 9-children #1 | Father | 82% | 7 | 2 | 0.5 | 0.09 |
| 9-children #2 | Father | 88% | 2 | 7 | $1 \times 10^{-5}$ | 0.09 |
| 9-children #3 | Father | 73% | 5 | 4 | 0.2 | 0.5 |
| 9-children #4 | Father | 71% | 3 | 6 | 0.02 | 0.25 |
| NC0 family | Mother | 91% | 1 | 1 | 0.17 | 0.75 |
| Combined | All | - | 37 | 45 | $3.6 \times 10^{-17}$ | 0.21 |

### Mutation transmission in families in Simons Simplex Collection

The Simons Simplex Collection contains whole-genome sequencing (WGS) data for 7,736 blood samples from 1,934 quartet families, each composed of a mother, a father, a child affected by autism spectrum disorder (ASD), and an unaffected sibling control^16^. We leveraged that data to discover post-zygotic developmental mutations marking the dominant lineage in parents and analyzed the mutations’ transmission to children. Developmental mutations were discovered by finding loci with evidence for three haplotypes. Namely, for a given locus every cell has two germline haplotypes A and B. When a locus has a developmental mutation (e.g., on haplotype A) then mutated cells will have a mutated haplotype A (A^m^) and an unaffected haplotype B. Bulk blood will consist of mutated and non-mutated cells and will, thus, have three haplotypes: A, A^m^, and B. We developed an algorithm that considered pairs of nearby (within 100 bps) single nucleotide variants detected in the blood such that overlapping reads can phase the somatic with germline variants and indicate the existence of three haplotypes (**Fig. 2A**). In such a pair of variants, one is an inhered heterozygous variant that serves as an anchor to phase the second one – an early developmental mutation – to haplotypes. We only considered developmental mutations at high frequency to ensure that they mark the dominant lineage.

To calibrate this approach and access its accuracy, we first applied it to mutation discovery in children in the quads (**Fig. 2B**). A true early post-zygotic mutation is likely to be unique and is unlikely to match variants in the corresponding parents. Thus, we considered a phased early mutation as a false positive if it had a matching variant call in one of the corresponding parents. We required VAF between 35% and 45% (corresponding to a cell frequency of 70-90%, since the dominant lineage should be in the majority of blood cells), and defined criteria on the number of reads supporting the minor (out of three) haplotype to eliminate most of the false positives. We arrived at a set of 35 early post-zygotic mutations in children with just 6 (17%) false positives (**Fig. 2C; Supplementary Table 3**). Of the remaining 29 early mutations in children, 25 (86%) have been previously identified as *de novo* using the standard approach by An *et al*.^16^ indicating that early mutations we discover are hardly distinguishable from *de novo* mutations.

We then analyzed mutations in parents, assuming the same false positive rate found in children, as WGS data for parents and children had similar coverage and other sequencing characteristics. We discovered a comparable number of 40 post-zygotic mutations in 40 parents of 40 families (**Fig. 2D; Supplementary Table 3**). Since every family in SSC has just 2 children, analysis of mutation transmission in each family lacks statistical power. We, therefore, analyzed the distribution of the 40 families by the outcomes of mutation transmission to their children, which were: transmission to none of the children, transmission to only one child, and transmission to both children (**Fig. 2E**). In this analysis, we considered four scenarios for the contribution of the dominant lineage to gonads and therefore to children: (i) frequency is the same as in blood (i.e., gonads and soma have similar lineage composition); (ii) frequency is 50% (i.e., equal contribution of dominant and recessive lineages to gonads); (iii) frequency is 100% (i.e., only dominant lineage is present in the gonads and passed to the next generation); (iv) frequency is random (**Supplementary Fig. 2**). For each scenario we derived the expected distribution of mutation transmission to none, one, or both children and compared it to the observed distribution (**Fig. 2E & Supplementary Fig. 3**). We could confidently reject the scenario of the same frequency of the dominant lineage in gonads and blood (*p*-value = 0.03) supporting such an inference made above. We could also confidently reject the scenario that only dominant lineage is present in the gonads (*p*-value = 10^-4^) (**Supplementary Fig. 3B**), supporting the inference made above for NC0 that the gonads are derived from at least two founder cells each descending from one blastomere likely arising from first zygotic cleavage. At the same time, the observed distribution was consistent with the equal contribution of dominant and recessive lineages to the gonads (*p*-value = 0.62) supporting the same observation made above for NC0 and large families (**Table 1**). The scenario of random distribution of lineages in the gonads is also consistent with the observations (*p*-value = 0.41). Yet note, that even in this scenario almost always the germline would consist of at least two cells – one from the dominant lineage and one from the recessive.

**Figure 3.**
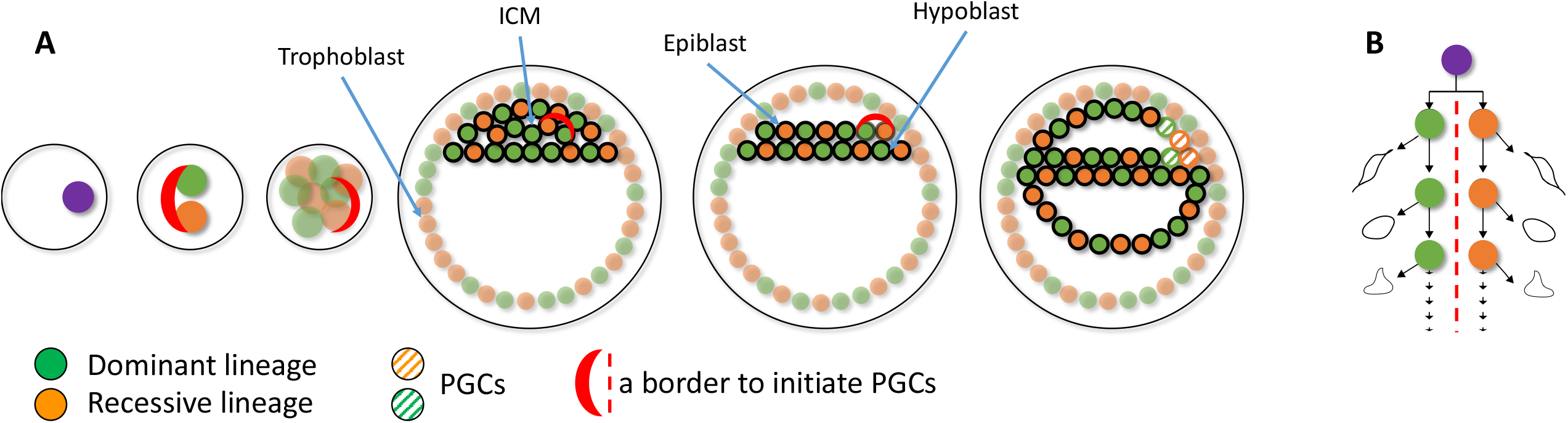
A possible model explaining differential lineage allocation in soma and germline. **A)** The first two blastomeres each founding the dominant and recessive lineage establish an invariant lineage border that is maintained through subsequent development perhaps through asymmetric division of the blastomeres (see B). After development of the epiblast, the border initiates the differentiation of the surrounding cells into PGCs, which results in roughly the same number of cells from the dominant and recessive lineages in the germline. **B)** The first blastomers may divide asymmetrically in the sense that only one of the two progenies stays at the border.

## Discussion

In the described study we leveraged three different approaches to study transmission of early mutations from parents to the next generation. The approaches included (i) direct reconstruction of developmental lineages in a parent and tracing their transmission to the progeny, (ii) identification of early mutations in parents of a collection of 1,934 familial quads, and (iii) analysis of early mutations in parents discovered as shared *de novo* mutations across multiple children. All approaches consistently revealed that both dominant and recessive developmental lineages in parental soma contribute to the germline, which consequently has at least two FGCs. Similarly, all the approaches consistently revealed differential frequencies of the dominant lineage in the germline and blood, suggesting a fundamental difference in lineage allocation between the soma and the germline. Moreover, all the approaches consistently suggest balanced (i.e. 50:50) allocation of dominant and recessive lineage to the germline, and we hypothesize that such an allocation is a common phenomenon in humans. This hypothesis is supported by direct measurement of frequencies of dominant lineage in the testis of individual PD28690 in a previous study^7^.

We and others previously hypothesized that asymmetric contribution of early lineages to soma could be the result of one of the first two blastomeres committing mostly to the extraembryonic (trophectoderm) lineages, with the other blastomere mostly committing to inner cell mass (ICM), becoming the dominant lineage in the embryo^5^. A recent imaging study of the developing human embryo supported the asymmetric contribution of the first two blastomeres to the ICM but suggested that the asymmetry arises from stochasticity in internalizing just a few, 1 to 4, cells to the ICM at the 8-to-16-cell stage^17^. If allocation of PGCs (that develop from the ICM) is also stochastic, then one could expect roughly the same bias in lineage contribution for the germline as for the soma. However, the results presented above (i.e., different frequencies of early lineage in germline and blood) contradict this expectation. This may imply that allocation of PGCs may not be stochastic and may depend on yet unknown factors, e.g., relative position in the ICM of the cells from dominant and recessive lineages, which eventually results in a different frequency of allocation of the lineages to soma and germline. In such scenario, similar contribution of dominant and recessive lineages to the germline may imply that PGCs are initiated at the border between one “dominant” and one “recessive” cell assemblies. Such a border may have roughly the same number of surrounding cells form dominant and recessive lineage committing to PGCs. This scenario could be consistent with the proposed dual origin of PGCs from amnion and epiblast in humans^18-21^. It is intriguing to speculate that there could be a molecular mechanism for maintaining such a border interface from the first zygotic cleavage until PGC specification perhaps trough asymmetric division of the blastomeres (**Fig. 3**).

Larger studies of mutation transmission in multi-children families combined with analysis of somatic and germline lineages in parents will likely shed light on this issue. More broadly, leveraging naturally occurring somatic mutations is a powerful approach to study human development, which will inform not only about the formation of the germline but also about other developmental steps prior and after gastrulation. Finally, these findings highlight the unique nature of early post-zygotic mutations, which is their ability to be transmitted to progeny and therefore contribute to the evolution of the human genome.

## Methods

### Somatic mosaicism and reconstruction of a cell lineage tree in the mother

In our previous study^5^, we reconstructed a cell lineage tree in early development by leveraging early post-zygotic mosaic mutations that we identified in the mother (NC0). In brief, we obtained whole genome sequencing data for 25 clonal induced pluripotent stem cell (iPSC) lines derived from skin fibroblasts (average coverage of 35x), as well as for bulk blood, saliva, and urine samples (average coverage of 238x) from NC0. Using the all-2-all exhaustive comparison^22^ of all sequenced iPSC lines, we discovered mosaic mutations which arose in development and accumulated in cell lineages. Based on the sharing of discovered early mutations and their estimated cell frequencies in bulk tissues, we reconstructed an early developmental cell ancestry tree starting from the likely first zygotic cleavage.

### Collection of samples and whole genome sequencing

Informed consent was obtained from each participant enrolled in the study according to the regulations of the Institutional Review Board and Yale Center for Clinical Investigation at Yale University. The participant agreed to data sharing of genomic de-identified data using controlled data access. About 14 mL of whole blood was collected for each individual using BD Vacutainer ACD tubes. DNA was extracted using the Maxwell® RSC Blood DNA Kit (Promega) in accordance with the manufacturer’s instructions. Whole Genome Sequencing was conducted to 30x coverage at BGI Americas Corporation for NC0-1 and NC0-2 and at the Yale Center for Genome Analysis for NC0-f. Measurements of DNA quantity by Qubit, and quality, by Agarose Gel Electrophoresis, were performed before and after the PCR-free library preparations.

### Calling constitutional/inherited germline variations in children and the father of NC0 family

The blood of two children of NC0 (NC0-1 and NC0-2) and father (NC0-f) was sequenced to 36x and 48x coverage on average, respectively. We aligned the raw sequences against the human reference genome hg19. The duplicated reads were marked and removed by Picard (http://broadinstitute.github.io/picard), and local realignment and base quality score recalibration were performed with GATK^23^. We used GATK HaplotypeCaller version 4.2.6 to call constitutional/inherited germline variants, resulting in 4,057,075 variants passing GATK filters with VQSR (Variant Quality Score Recalibration) in the father, and 4,231,719 and 4,167,927 in the two daughters.

### Identification of transmitted early mutations in NC0

Out of all discovered post-zygotic mosaic mutations in the mother (18,419; see above), we retained only the mutations matching constitutional/inherited germline variations found in each daughter (see above) as candidates for transmitted early post-zygotic mutations. This resulted in 27 candidates, of which 18 were excluded as they were present in almost 100% (i.e., at 50% VAF) of the father’s blood cells, thereby indicating they were inherited from the father **(Supplementary Table 2)**. Moreover, those excluded mutations were not shared by any other sampled lineage and lacked supporting reads in bulk tissues (blood, saliva, and urine) in the mother, suggesting they occurred very late in development, are private to the specific somatic lineages where they were found and not relevant to the maternal germline lineage. Consequently, we identified 2 and 7 early post-zygotic mutations in the mother transmitted to each of the offspring. The constitutional nature of these mutations in the daughters was evidenced by their VAFs of around 50% in the blood **(Fig. 1D and Supplementary Table 1)**, which indicated that they were also present in germline lineages in the mother, thus had occurred prior to the differentiation and proliferation of primordial germ cells (PGCs) (**Fig. 1C**).

### Extraction of *de novo* candidates through the whole genome comparison within the parents-offspring trio

We searched *de novo* mutations within the trio by comparing the genomes of two children to the maternal and paternal genomes. To mitigate the potential effects of coverage bias, we down-sampled the maternal genome from its original average coverage of 238x to 48x, which is equivalent to the average coverage of the paternal genome. We used Mutect2^23^ and Strelka2^24^ to call mutations in each child against their parental genomes and considered only consensus “PASS” calls made by both callers. Calls were required to have a VAF of between 30% and 70% in the child and a minimum depth of 20 reads in the child and both parents. We also excluded sites within known segmental duplication and simple tandem repeat regions, as defined by the UCSC (https://genome.ucsc.edu/cgi-bin/hgTables). For each child, we defined mutations as *de novo* if they satisfied such criteria and were called relative to both parental genomes. This yielded 84 *de novo* mutation candidates in one child and 94 in the other. We observed that only 4 out of 9 early post-zygotic mutations in the mother, which were transmitted to the offspring, were also identified as *de novo* mutations in children using this trio whole genome comparison strategy **(Supplementary Table 1)**. There was only one shared *de novo* candidate variant between daughters. Given that lineage reconstruction in mother suggest that transmitted mutations to daughters can’t be shared (transmitted lineages as originating from the likely first zygotic cleave share not common mutations), this variant likely originated in father. Alternatively, it could be a systematic false positive.

Furthermore, we additionally applied the parental presence filter as in other trio studies^4,14,15^, where *de novo* mutations were required to have a maximum depth of 1 read and a maximum allele frequency of 5% for alternative alleles in both parents. Only 1 out of the 9 early post-zygotic mutations in the mother transmitted to children was retained as *de novo*, with a total of 79 in one child and 82 in the other. This demonstrates that the number of early developmental mutations transmitted to offspring, which likely occurred before the specification of PGCs (PGCS), may have been underestimated with the standard trio approach^4,14,15^. This is due to the exclusion of variations with evidence of presence in parents, which removes a large proportion of transmitted mutations that arose in pre-PGCS lineages. Some of these *de novo* mutations may be recovered with the haplotype sharing by siblings’ approach^10,11^, but the approach still misses *de novo* mutations in the germline that are carried by all or none of siblings who share the same parental haplotype at the locus^10^.

### Analyzing large families from Iceland pedigree study

We utilized data presented in Jonsson *et al*.^10^ in figure 2 and supplementary figures 1 and 2. For each parent we defined lineages by branches from the ancestral states defined by Jonsson *et al*. There were 2 to 3 of such lineages/branches for parents in the family. Since no lineage tree were reconstructed in each parent, no clear mutation marker(s) for the lineages were defined. Therefore, we estimated frequencies of lineages in the blood from the frequencies of transmitted mutations in the following ways. Mutations with higher frequencies mark earlier originating lineages, so, for each lineage, we first identified a primary marker mutation as a mutation with the highest VAF in the blood. We then defined a set of lineage markers as mutations with frequencies different by less than 10% VAF from the frequency of the primary marker. We then averaged VAFs of the defined marker set and estimated cell frequencies of the corresponding lineages as a double of the average VAF.

We used 7 parents with an obvious dominant lineage in the soma (i.e., at least 70% of cells in the blood). Frequency of the recessive lineage was defined as a complement to the frequency of the dominant lineage. Accordingly, counts of children derived from the transmitted recessive lineage were calculated as the number of children without transmitted dominant lineage. When testing for a consistency with the count of children with the transmitted lineage and the frequency of the lineage in the blood we used one-tailed binomial test for less or equal count of children with the dominant lineage.

### Analyzing transmission of early mutations across generations in the Simons Simplex Collection

We downloaded CRAM files (aligned to the human reference genome GRCh38) and VCF files from SFARI Base (https://www.sfari.org/resource/sfari-base). To select inherited SNPs for phasing with mutation candidates (**Fig. 2A**), we searched heterozygous single nucleotide variant calls from VCF files. We retained only calls that passed GATK filters, were in accessible genomic regions according to the 1000 Genomes Project’s mappablility mask (bases marked as “P”) and had a population allele frequency of more than 10^-3^ in gnomAD database (https://gnomad.broadinstitute.org). Mutation candidates were selected from single nucleotide variant calls within 100 bps upstream and downstream of the selected inherited SNPs. To retain the most likely early mutations we required the calls to be in genomic regions with the “P” mask, pass GATK filters, have a population allele frequency of less than 10^-5^, and be outside the known repeat regions defined by the UCSC table browser. We then used a phasing tool in MosaicForecast package^25^ to phase two nearby SNVs (inherited SNP and candidate mutation). We retained candidate mutations that were phased to 3 haplotypes, had at least 3 supporting reads for the minor haplotype, and had a VAF of more than 35% for further consideration.

For the final sets of phased early post-zygotic mutations we required that the minor haplotype had 4 to 6 supporting reads for VAF [35%,40%] and 5 or 6 supporting reads for VAF [40%,45%]. This was based on the analysis for children, for which we considered a phased early mutation as a false positive if there was a matching variant call in parents, i.e., the mutation was an inherited SNP. We additionally removed phased clustered within 100 bps mutations. For the final set of 35 early developmental mutations in children the false positive rate was estimated to be 17% (**Fig. 2C**). Except for these false positives, none other variants in the set had any supporting reads in the parents, suggesting the calls are genuine early developmental mutations. Because WGS data for parents and children had similar coverage and other sequencing characteristics, we assume the same false positive rate for the final set of phased early mutations in parents (**Fig. 2D**).

We calculated transmission probabilities for early mutations in a parent to none, one, or two children for four possible scenarios based on their expected frequencies in germ cells of the parent, i.e., based on the contribution of the dominant lineage to germline development (**Supplementary Fig. 2B**). For the scenario where the frequency of the dominant lineage in gonads is same as in blood, we first calculated transmission probabilities for each mutations and then averaged the probabilities across the entire set of mutations. To derive the expected distribution of mutation transmission, we applied the 17% false positive rate for inherited variants (**Fig. 2E** and **Supplementary Fig. 3**), as estimated in the analysis for children. We considered a candidate early mutation in parents as a transmitted one if it had a matching variant call that passes GATK filters in one or two corresponding children. To test the statistical difference between expected and observed transmission distributions, we used *χ*^2^ test with 2 degrees of freedom.

## Data availability

All primary data have been uploaded to the NIMH Data Archive (NDA) under collection #2961, at url: https://nda.nih.gov/edit_collection.html?id=2961. WGS and phenotype data for the SSC collection are hosted by the Simons Foundation for Autism Research Initiative (SFARI) and are available for approved researchers at SFARI Base (https://base.sfari.org/).

## Ethics statement

Human subjects were recruited through several research projects at the Yale Child Study Center. Written informed consent was obtained from each participant enrolled in the study, and all research was approved by the Yale University Institutional Review Board (HIC# 1104008337) and Yale Center for Clinical Investigation at Yale University. Analysis of SSC data was conducted under IRB 23-012592 approved by Mayo Clinic IRB.

## Acknowledgments

We are grateful to member of NC0 family that participated in this study by donating tissue and/or blood samples. We are grateful to members of families in SSC that donated blood samples for WGS and to Simons Foundation that provided access to the data (approved project 2343.4). This work was funded by the NIH Common Fund SMaHT program (grants UG3 NS132128 and UG3 NS132146) and by the Simons Foundation (grant 399558). Y.J. was also supported by Basic Science Research Program through the National Research Foundation of Korea (NRF) funded by the Ministry of Education (grant number 2022R1A6A3A03055692).

## Author contributions

F.M.V. and A.A. conceived the study. A.A. and F.M.V. supervised the study. L.T. and A.S. collected samples and performed experiments. Y.J. and A.A. performed computational data analyses and prepared display items. F.M.V., A.A., and Y.J. drafted the initial text of the manuscript.

